# Inferring parameters of the distribution of fitness effects of new mutations when beneficial mutations are strongly advantageous and rare

**DOI:** 10.1101/855411

**Authors:** Tom R. Booker

## Abstract

Characterising the distribution of fitness effects (DFE) for new mutations is central in evolutionary genetics. Analysis of molecular data under the McDonald-Kreitman test has suggested that adaptive substitutions make a substantial contribution to between-species divergence. Methods have been proposed to estimate the parameters of the distribution of fitness effects for positively selected mutations from the unfolded site frequency spectrum (uSFS). However, when beneficial mutations are strongly selected and rare, they may make little contribution to standing variation and will thus be difficult to detect from the uSFS. In this study, I analyse uSFS data from simulated populations subject to advantageous mutations with effects on fitness ranging from mildly to strongly beneficial. When advantageous mutations are strongly selected and rare, there are very few segregating in populations at any one time. Fitting the uSFS in such cases leads to underestimates of the strength of positive selection and may lead researchers to false conclusions regarding the relative contribution adaptive mutations make to molecular evolution. Fortunately, the parameters for the distribution of fitness effects for harmful mutations are estimated with high accuracy and precision. The results from this study suggest that the parameters of positively selected mutations obtained by analysis of the uSFS should be treated with caution and that variability at linked sites should be used in conjunction with standing variability to estimate parameters of the distribution of fitness effects in the future.

## Introduction

Characterising the distribution of fitness effects for beneficial mutations is central in evolutionary biology. The rate and fitness effects of advantageous mutations may determine important evolutionary processes such as how variation in quantitative traits is maintained (Hill, 2010), the evolution of sex and recombination (Otto, 2009) and the dynamics of evolutionary rescue in changing environments (Orr & Unckless, 2014). However, despite its central role in evolution, relatively little is known about the distribution of fitness effects (DFE) for advantageous mutations in natural populations. The DFE for advantageous mutations can be estimated from data obtained via targeted mutation or from mutation accumulation experiments (e.g. Bank, Hietpas, Wong, Bolon, & Jensen, 2014; Böndel et al., 2019; reviewed in Bailey & Bataillon, 2016), but such efforts may be limited to laboratory systems. Alternatively, estimates of the DFE can be obtained for natural systems using population genetic methods.

When natural selection is effective, beneficial alleles are promoted to eventual fixation while deleterious variants are maintained at low frequencies. Migration, mutation, selection and genetic drift interact to shape the distribution of allele frequencies in a population (Wright, 1937). Parameters of the DFE for both advantageous and deleterious mutations can be estimated by modelling population genomic data, specifically the site frequency spectrum (SFS). The SFS is the distribution of allele frequencies present in a sample of individuals drawn from a population. By contrasting the SFS for a class of sites expected to be subject to selection with that of a neutral comparator, one can estimate the parameters of the DFE if selected mutations are segregating in the population of interest (reviewed in Eyre-Walker & Keightley, 2007). Typically, the DFE for nonsynonymous sites in protein coding genes is estimated using synonymous sites as the neutral comparator. Several methods have been proposed that estimate the DFE for deleterious mutations from the SFS under the assumption that beneficial mutations contribute little to standing genetic variation (e.g. Barton & Zeng, 2018; Boyko et al., 2008; Keightley & Eyre-Walker, 2007; Tataru, Mollion, Glemin, & Bataillon, 2017).

The DFE for deleterious mutations can be used when estimating *α*, the proportion of between-species divergence attributable to adaptive evolution (Eyre-Walker & Keightley, 2009). *α* can be estimated by rearranging the terms of the McDonald-Kreitman test (MK-test), which assesses the extent of positive selection. Under strong purifying selection, the ratio of divergence at nonsynonymous sites (*d*_*N*_) to that of synonymous sites (*d*_*S*_) should be exactly equal to the ratio of nucleotide diversity at nonsynonymous (*π*_*N*_) and synonymous sites (*π*_*S*_)(McDonald & Kreitman, 1991). Adaptive evolution of protein sequences may contribute to *d*_*N*_ such that *d*_*N*_/*d*_*S*_ > *π*_*N*_/*π*_*S*_. Charlesworth (1994) suggested rearranging the terms of the MK-test to estimate the excess *d*_*N*_ due to positive selection (*α*) as

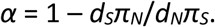

Slightly deleterious alleles may contribute to both standing genetic variation and between-species divergence, estimates of *α* may therefore be refined by subtracting the contribution that deleterious alleles make to both polymorphism and divergence and this can be calculated using the DFE for harmful mutations (Eyre-Walker & Keightley, 2009). Application of such methods to natural populations suggest that *α* is of the order of 0.5 in a large variety of animal taxa (Galtier, 2016). However, if adaptive evolution is as frequent as MK-test analyses suggest, the assumption that advantageous alleles contribute little to standing variation may be violated and ignoring them could lead to biased estimates of the DFE (Tataru et al., 2017).

When advantageous alleles contribute to standing variation, parameters of the DFE for both deleterious and beneficial mutations can be estimated from the SFS (Schneider et al., 2011; Tataru et al., 2017). When data from an outgroup species are available, variable sites within a focal species can be polarised as either ancestral or derived and the *unfolded* SFS (uSFS) can be obtained. Inference of ancestral/derived states is, however, potentially error-prone (Keightley & Jackson, 2018). The uSFS is a vector of length 2*n*, where *n* is the number of haploid genome copies sampled. The *i*^th^ entry of the uSFS is the count of derived alleles observed at a frequency *i* in the sample. Note that when outgroup data are not available, alleles cannot be polarised and the distribution of minor allele frequencies (known as the *folded* SFS) is analysed. There is limited power to detect positive selection from the SFS, so the DFE for beneficial mutations is often modelled as a discrete class of mutational effects, with one parameter specifying the fitness effects of beneficial mutations, *γ*_a_ = 2*N*_*e*_*s*_*a*_ where *N*_*e*_ is the effective population size and *s*_*a*_ is the positive selection coefficient in homozygotes, and another specifying the proportion of new mutations that are advantageous, *p*_*a*_. Estimates of *γ*_*a*_ and *p*_*a*_ for nonsynonymous sites have only been obtained a handful of species, and these are summarised in Table 1. The positive selection parameter estimates that have been obtained for mice and *Drosophila* are fairly similar (Table 1). Note that the estimates for humans obtained by Castellano et al, (2019) did not provide a significantly greater fit to the observed data than did a model with no positive selection. Furthermore, Castellano et al, (2019) estimated the parameters for numerous great ape species, the parameters shown for humans are representative of the estimates for all taxa they analysed.

**Table 1.** Estimates of the parameters of positive selection obtained from the uSFS for nonsynonymous sites.

| Common name | Scientific name | $\gamma_a$ | $p_a$ | Reference | Method used<sup>¶</sup> |
| --- | --- | --- | --- | --- | --- |
| House mouse | *Mus musculus castaneus* | 14.5 | 0.0030 | Booker & Keightley, (2018) | DFE-alpha |
| Fruit fly | *Drosophila melanogaster* | 23.0 | 0.0045 | Keightley et al, (2016) | DFE-alpha |
| Humans | *Homo sapiens* | 0.0064<sup>†</sup> | 0.000025 | Castellano et al, (2019) | polyDFE |

<sup>¶</sup> - DFE-alpha implements the analysis methods described by Schneider et al., (2011), polyDFE implements the methods described by Tataru et al., (2017)
† - Castellano et al., (2019) estimated the mean fitness effect for an exponential distribution of advantageous mutational effects.

Depending on the rate and fitness effects of beneficial mutations, different aspects of population genomic data may be more or less informative for estimating the parameters of positive selection. As beneficial mutations spread through populations, they may carry linked neutral variants to high frequency, causing selective sweeps (Barton, 2000). On the other hand, if advantageous mutations have mild fitness effects, they may take a long time to reach fixation and make a substantial contribution to standing genetic variation. Because of this, uSFS data and polymorphism data at linked sites may both be informative for understanding the parameters of positive selection. For example, Campos et al., (2017) used a model of selective sweeps to analyse the negative correlation observed between *d*_*N*_ and *π*_*S*_ in *Drosophila melanogaster* and estimated *γ*_a_ = 250 and *p*_*a*_ = 2.2 × 10^−4^, but this method assumes a constant population size. An analysis of the uSFS from the same dataset that modelled of population size change yielded estimates of *γ*_a_ = 23 and *p*_*a*_ = 0.0045 for nonsynonymous sites (Keightley et al., 2016). The sharp contrast between the two studies’ estimates of the positive selection parameters may due to different assumptions but could potentially be explained if the DFE for advantageous mutations in *D. melanogaster* is bimodal. If this were so, the different methods (i.e. sweep models versus uSFS analysis) may be capturing distinct aspects of the DFE for advantageous mutations, or it could be that both models are highly unidentifiable. The handful of studies that have attempted to estimate *γ*_a_ and *p*_*a*_ from the uSFS have yielded similar estimates of positive selection (Table 1), which may indicate commonalities in the DFE for beneficial mutations across taxa. On the other hand, uSFS analyses may have only found evidence for mildly beneficial mutations because the approach is only powered to detect weakly beneficial mutations. Indeed, verbal arguments have suggested that rare strongly selected advantageous mutations, which may contribute little to standing variation, will be undetectable by analysis of the uSFS (Booker & Keightley, 2018; Campos et al., 2017).

The studies describing the two most recently proposed methods for estimating the DFE for beneficial mutations from the uSFS (Schneider et al., 2011; Tataru et al., 2017) performed extensive simulations, but did not test cases of rare advantageous mutations with strong effects on fitness. Testing this case is important, as studies that have analysed patterns of putatively neutral genetic diversity across the genome have indicated that the DFE for advantageous mutations contains strongly beneficial mutations in a variety of taxa (Booker & Keightley, 2018; Campos et al., 2017; Elyashiv et al., 2016; Nam et al., 2017; Uricchio et al., 2019). Note that Tataru et al., (2017) did simulate a population subject to frequent strongly beneficial mutations (*γ*_a_ = 800 and *p*_*a*_ = 0.02), but the parameter combination they tested may not be biologically relevant as the proportion of adaptive substitutions it yielded was far higher than is typically estimated from real data (*α =* 0.99). The limited parameter ranges tested in the simulations performed by Schneider et al., (2011) and Tataru et al., (2017) leave a critical gap in our knowledge as to how uSFS based methods perform when advantageous mutations are strongly selected and infrequent.

In this study, I use simulated datasets to fill this gap and examine how uSFS-based analyses perform when beneficial mutations are strongly selected and rare. I simulate populations subject to a range of positive selection parameters, including cases similar to those modelled by Tataru et al., (2017) and cases where beneficial mutations are strongly selected but infrequent. It has been pointed out that estimating selection parameters by modelling within species polymorphism along with between-species divergence makes the assumption that the DFE has remained invariant since the ingroup and outgroup began to diverge (Tataru et al., 2017). By analysing only the polymorphism data, one can potentially avoid that problematic assumption. Using the state-of-the-art package *polyDFE* v2.0 (Tataru & Bataillon, 2019), I analyse the uSFS data and estimate selection parameters for all simulated datasets with or without divergence. The results from this study suggest that, when beneficial mutations are strongly selected and rare, analysis of the uSFS results in spurious parameter estimates and the proportion of adaptive substitutions may be poorly estimated.

## Methods

### Population genomic simulations

I tested the hypothesis that the parameters of infrequent, strongly beneficial mutations are difficult to estimate by analysis of the uSFS using simulated datasets. Wright-Fisher populations of *N*_*e*_ = 10,000 diploid individuals were simulated using the forward-in-time package *SLiM* (v3.2; Haller & Messer, 2019). Simulated chromosomes consisted of seven gene models, each separated by 8,100bp of neutrally evolving sequence. The gene models consisted of five 300bp exons separated by 100bp neutrally evolving introns. The gene models were based on those used by Campos & Charlesworth, (2019), but unlike that study, I did not model the untranslated regions of genes. Nonsynonymous sites were modelled by drawing the fitness effects for 2/3rds of mutations in exons from a distribution of fitness effects (DFE), while the remaining 1/3 were strictly neutral and used to model synonymous sites. The fitness effects of nonsynonymous mutations were beneficial with probability *p*_*a*_ or deleterious with probability 1 – *p*_*a*_. Beneficial mutations had a fixed selection coefficient of *γ*_*a*_ = *2N*_*e*_*s*_*a*_. The fitness effects of deleterious mutations were drawn from a gamma distribution with a mean of *γ*_*d*_ = 2*N*_*e*_*s*_*d*_ = −2,000 and a shape parameter of *ß =* 0.3 (*s*_*d*_ being the negative selection coefficient in homozygotes). The gamma distribution of deleterious mutational effects was used for all simulated datasets and was based on results for nonsynonymous sites in *Drosophila melanogaster* (Loewe & Charlesworth, 2006). Uniform rates of mutation (*μ*) and recombination (*r*) were set to 2.5 × 10^−7^ (giving *4N*_*e*_*r* = *4N*_*e*_*µ* = 0.01). Note that *μ* and *r* are far higher than is biologically realistic for most eukaryotes, I scaled up these rates to model a population with a large *N*_*e*_ using simulations of 10,000 individuals. Across simulations I varied the *γ*_*a*_ and *p*_*a*_ parameters and performed 2,000 replicates for each combination of parameters. Thus, I simulated a dataset of 21Mbp of coding sequence for each combination of *γ*_*a*_ and *p*_*a*_ tested.

In this study, I assumed a discrete class of beneficial mutational effects rather than a continuous distribution, which is likely unrealistic for most organisms. Theoretical arguments have been proposed that the DFE for beneficial mutations that go to fixation should be exponential (Orr, 2003). However, the studies that have estimated the DFE for beneficial mutations from population genetic data have often modelled discrete classes of effects (Campos et al., 2017; Elyashiv et al., 2016; Keightley et al., 2016; Uricchio et al., 2019). I chose to model discrete selection coefficients in the simulated datasets in order to better understand the limitations of the methods rather than to accurately model the DFE for beneficial mutations.

To model the accumulation of nucleotide substitutions after the split of a focal population with an outgroup, I recorded all substitutions that occurred in the simulations. Campos & Charlesworth, (2019) analysed simulations very similar to those that I performed in this study and showed that populations subject to beneficial mutations with *γ*_*a*_ = 250 and *p*_*a*_ = 0.0002 took 14*N*_*e*_ generations to reach mutation-selection-drift equilibrium. In this study I modelled a range of positive selection parameters, so to ensure that my simulations reached equilibrium I performed 85,000 (34*N*_*e*_) generations of burn-in before substitutions were scored. The expected number of neutral nucleotide substitutions that accumulate per site in *T* generations is *d*_*Neutral*_ = *Tµ*. The point mutation rate in my simulations was set to *µ* = 2.5 × 10^−7^per site per generation, so I ran the simulations for 200,000 generations beyond the end of the burn-in phase to model a neutral divergence of *d*_*Neutral*_ = 0.05. All variants present in the population sampled at a frequency of 1.0 were also scored as substitutions.

Using the 2,000 simulated datasets, I constructed 100 bootstraps by sampling with replacement. From each bootstrap sample, I collated variants and constructed the uSFS for synonymous and nonsynonymous sites for 20 diploid individuals.

### Analysis of simulation data

I calculated several summary statistics from the simulated datasets. Firstly, I calculated pairwise nucleotide diversity at synonymous sites (*π*_*s*_) and expressed it relative to the neutral expectation of *π*_*0*_ = 4*N*_*e*_*µ* = 0.01. Secondly, divergence at nonsynonymous sites for both advantageous (*dN*_*a*_) and deleterious mutations (*dN*_*d*_) was used to calculate the observed proportion of adaptive substitutions, *α*_*Obs*_ = *dN*_*a*_/(*dN*_*a*_ + *dN*_*d*_). Finally, I recorded the total number of beneficial mutations segregating in simulated populations, *S*_*Adv*_.

I estimated DFEs from simulated data by analysis of the uSFS using *polyDFE* (v2.0; Tataru & Bataillon, 2019). *polyDFE* fits an expression for the uSFS expected under a full DFE to data from putatively neutral and selected classes of sites and estimates parameters by maximum likelihood. For each set of positive selection parameters, simulated uSFS data were analysed under “Model B” in *polyDFE* (a gamma distribution of deleterious mutational effects plus a discrete class of advantageous mutations). Initial parameters for the maximisation were calculated from the data using the ‘-e’ option and the uSFS was analysed either with or without divergence using the “-w” option in *polyDFE*. Analysing the uSFS without divergence causes the selection parameters to be inferred from polymorphism data alone. For each replicate, I tested whether the inclusion of beneficial mutations in the DFE improved model fit using likelihood ratio tests between the best-fitting model and a model with *p*_*a*_ set to 0.0. Setting *p*_*a*_ = 0.0 means that positive selection does not influence the likelihood, so two fewer parameters are being estimated. Twice the difference in log-likelihood between the full DFE model and the model with *p*_*a*_ = 0.0 was tested against a *χ*^2^ distribution with 2 degrees of freedom. Likelihood surfaces were estimated by running *polyDFE* using a grid of fixed values for DFE parameters.

### Data Availability

All code and *SLiM* configuration files needed to reproduce the results shown in this study are available at https://github.com/TBooker/PositiveSelection_uSFS.

## Results

### Population genomic simulations

I performed simulations that modelled genes subject to mutation-selection-drift balance with fitness effects drawn from a distribution that incorporated both deleterious and advantageous mutations. The DFE for harmful mutations was constant, but I varied the fraction (*p*_*a*_) and fitness effects (*γ*_*a*_) of beneficial mutations across simulated datasets (Table 2). For each set of advantageous mutation parameters, 21Mbp of coding sequences was simulated, of which 14Mbp were nonsynonymous and 7Mbp were synonymous sites. Variants present in the simulated populations were used to construct the uSFS for a sample of 20 diploid individuals (Figure S1), a sample size which is fairly typical of current population genomic datasets (e.g. Castellano et al., 2019; Laenen et al., 2018; Williamson et al., 2014).

**Table 2.** Parameters of positive selection assumed in simulations and the proportion of *polyDFE* runs for which modelling positive selection gave a significantly better fit to the data.

| $\gamma_a$ | $p_a$ | $\gamma_a p_a$ | Proportion of likelihood ratio tests significant | | Proportion of analyses with gradient < 0.01 | |
| --- | --- | --- | --- | --- | --- | --- |
|  |  |  | With divergence | Without divergence | With divergence | Without divergence |
| 10 | 0.0001 | 0.001 | 0.02 | 0.07 | 0.11 | 0.71 |
| 50 |  | 0.005 | 0.98 | 0.86 | 0.10 | 0.77 |
| 100 |  | 0.01 | 0.98 | 0.02 | 0.03 | 0.58 |
| 500 |  | 0.05 | 1.00 | 0.39 | 0.00 | 0.99 |
| 1,000 |  | 0.10 | 1.00 | 1.00 | 0.00 | 0.71 |
| 10 | 0.001 | 0.01 | 0.99 | 0.96 | 0.15 | 0.71 |
| 50 |  | 0.05 | 1.00 | 1.00 | 0.06 | 0.98 |
| 100 |  | 0.10 | 1.00 | 1.00 | 0.00 | 0.97 |
| 500 |  | 0.50 | 1.00 | 1.00 | 0.00 | 0.94 |
| 1,000 |  | 1.00 | 1.00 | 1.00 | 0.00 | 0.71 |
| 10 | 0.01 | 0.10 | 1.00 | 1.00 | 0.03 | 0.80 |
| 50 |  | 0.50 | 1.00 | 1.00 | 0.02 | 0.99 |
| 100 |  | 1.00 | 1.00 | 1.00 | 0.02 | 0.95 |
| 500 |  | 5.00 | 1.00 | 1.00 | 0.00 | 0.72 |
| 1,000 |  | 10.0 | 1.00 | 1.00 | 0.00 | 0.41 |

Across simulations, the strength of selection acting on advantageous mutations ranged from *γ*_*a*_ = 10 to *γ*_*a*_ = 1,000. For a given *p*_*a*_ parameter, increasing the strength of selection increased the observed proportion of adaptive substitutions, *α*_*Obs*_ (Figure 1A). This is expected and is due to the monotonic increasing relationship between fixation probability and the strength of positive selection first described by Haldane (1927). Additionally, parameter combinations for which *γ*_*a*_*p*_*a*_ were equal had similar proportions of adaptive substitutions, for example compare *γ*_*a*_ = 10 and *p*_*a*_ = 0.01 to *γ*_*a*_ = 1,000 and *p*_*a*_ = 0.0001 (Figure 1A). This was also expected because the rate of adaptive substitutions is proportional to *γ*_*a*_*p*_*a*_. In some datasets, particularly when *p*_*a*_ = 0.01 and advantageous mutations were very strongly selected (i.e. *γ*_*a*_ ≥ 500), *α*_*Obs*_ exceeded 0.75, which is higher than is typically estimated from empirical data (Galtier, 2016), so these parameter combinations may not be biologically relevant.

**Figure 1.**
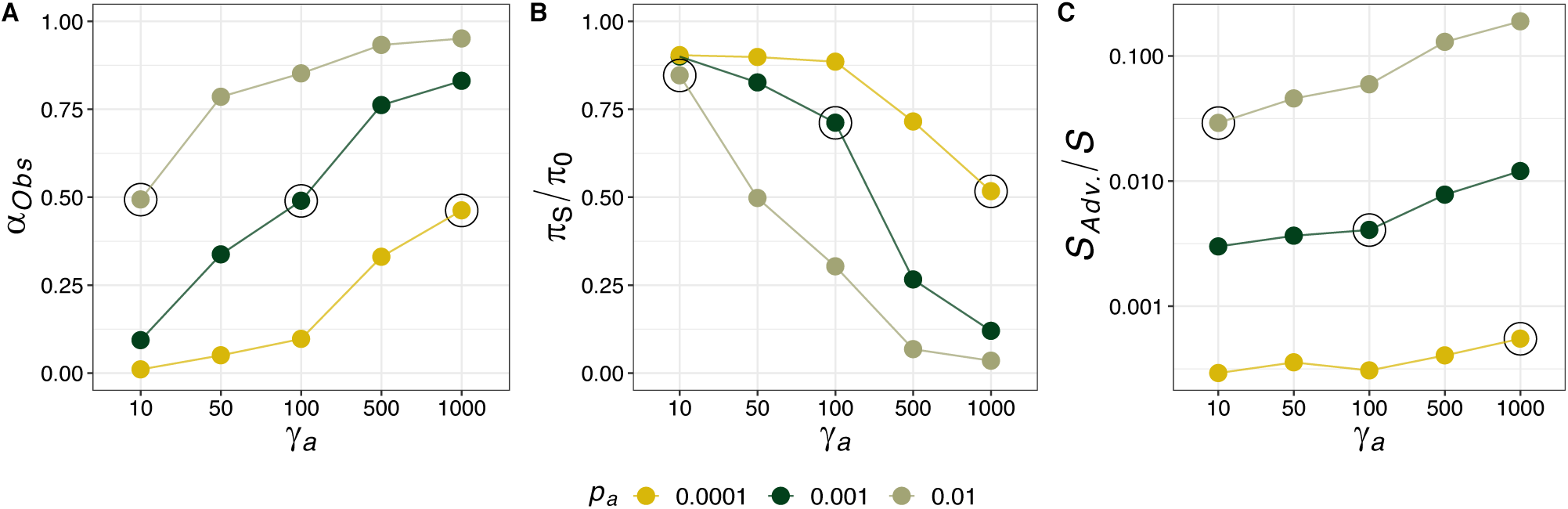
Population genetic summary statistics collated across all simulated genes. *α*_*Obs*_ is the observed proportion of substitutions fixed by positive selection. π_s_/π_0_ is genetic diversity relative to neutral expectation (*π*_*0*_ = 0.01). *S*_*Adv.*_/*S* is the proportion of segregating nonsynonymous sites that are advantageous in the simulated datasets.

The effects of selection at linked sites varied across simulated datasets. The DFE for deleterious mutations was kept constant across simulations, so the extent of background selection should be fairly similar across all parameter sets and thus variation in *π*_*S*_*/π*_*0*_ reflects the effects of selective sweeps. Under neutrality *π*_*S*_*/π*_*0*_ had an expected value of 1.0 and I found that selection at linked sites reduced nucleotide diversity below that expectation in all simulations (Figure 1B). Increasing the fitness effects or frequency of advantageous mutations had a strong effect on genetic diversity at synonymous sites, as shown by *π*_*S*_/*π*_*0*_ in Figure 1B. The highlighted points in Figure 1 indicate parameter combinations for which *γ*_*a*_*p*_*a*_ = 0.01. As expected, *α*_*Obs*_ for these three parameter sets was very similar (Figure 1A). Figure 1B shows that *π*_*S*_*/π*_*0*_ decreased across these three parameter combinations as the strength of positive selection increased. Finally, differences in *p*_*a*_ explained most of the variation in the proportion of segregating advantageous mutations (*S*_*Adv.*_*/S*) across simulated datasets, but *S*_*Adv.*_*/S*. also increased with the strength of positive selection (Figure 1C). On the basis of these results, it is clear that there will be lower power to estimate positive selection on the basis of standing variation when advantageous mutations are rare (i.e. *p*_*a*_ = 0.0001) than when they are comparatively frequent (i.e. *p*_*a*_ = 0.01).

### Analysis of the unfolded site frequency spectrum

Figure 2 shows the observed (bars) and expected (lines) distribution of derived allele frequencies for beneficial mutations segregating in simulated populations. The three panels of Figure 2 correspond to three parameter combinations for which *γ*_*a*_*p*_*a*_ = 0.01 (*γ*_*a*_ = 1,000 and *p*_*a*_ = 0.0001, *γ*_*a*_ = 100 and *p*_*a*_ = 0.001 and *γ*_*a*_ = 10 and *p*_*a*_ = 0.01). The lines in each of the panels of Figure 2 show the analytical expectation for the uSFS of advantageous mutations calculated using Equation 2 from Tataru et al., (2017). The analytical expectation closely matches the observed data for all three combinations (Figure 2). However, for a given value of *p*_*a*_, the analytical expectation for models with increasing fitness effects were very similar, which likely makes it difficult to distinguish them on the basis of polymorphism alone (Figure 2). For the three parameter sets shown in Figure 2, the overall contribution that advantageous alleles make to the uSFS for nonsynonymous sites is small relative to deleterious ones (Figure S1). Accurate estimation of positive selection parameters from the uSFS requires that the distribution of advantageous alleles can be distinguished from deleterious variants, so when *p*_*a*_ is small it seems likely that uSFS analyses will be unable to easily distinguish competing models.

**Figure 2.**
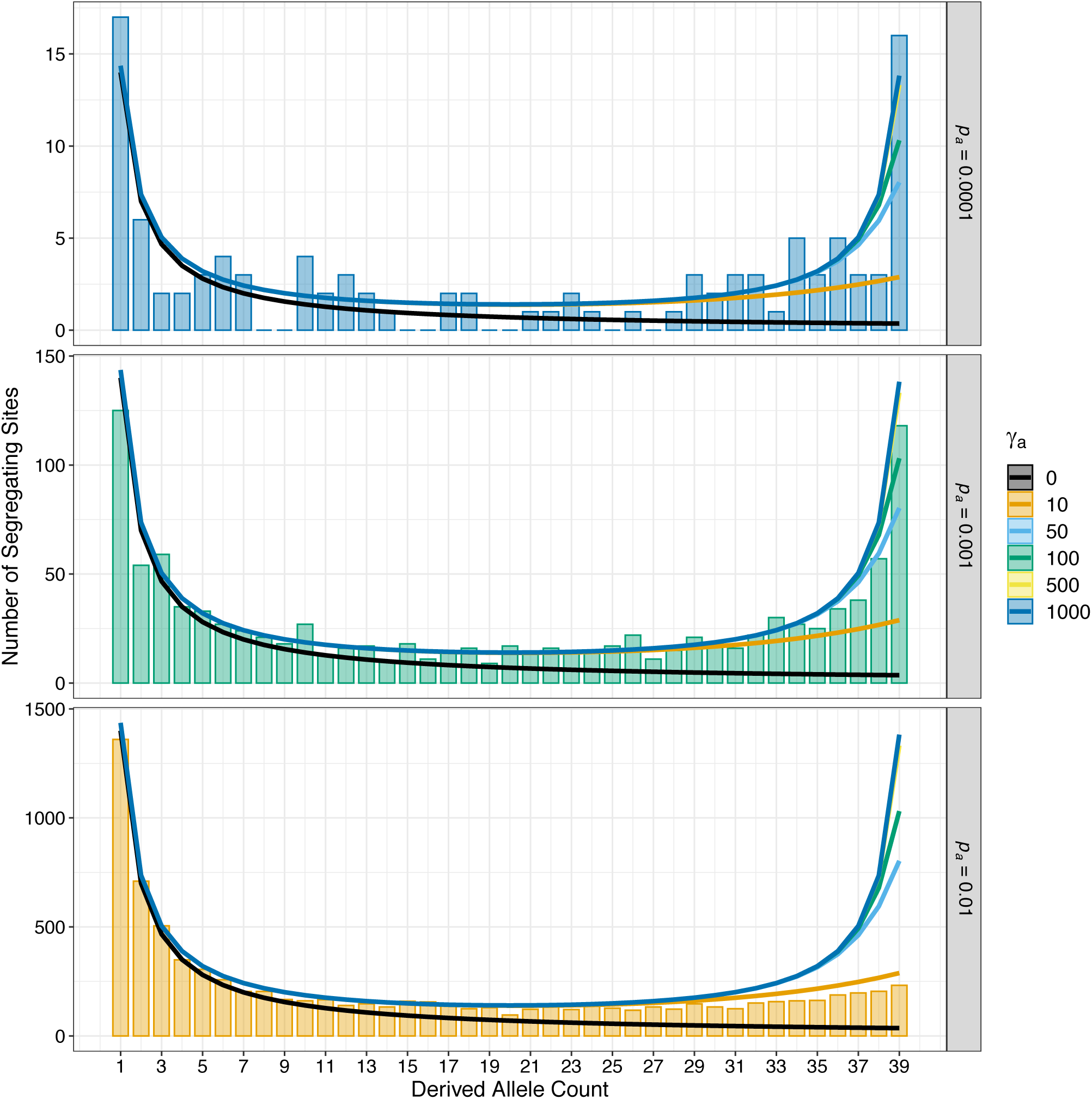
The uSFS for advantageous mutations under different combinations of positive selection parameters. The three bar charts show observed uSFS from simulations that model positive selection parameters that yield similar *α*. The lines in each panel show the expected frequency spectra for different strengths of beneficial mutations and were obtained using Equation 2 from Tataru et al., (2017).

When analysing a particular uSFS dataset in *polyDFE*, I either modelled the full DFE (i.e. a gamma distribution of deleterious mutations and a discrete class of advantageous mutational effects), or just a gamma DFE for harmful mutations (dDFE). I compared the two models using likelihood ratio tests, which tested the null hypothesis that the fit of the full DFE model is similar to that of a model containing only deleterious mutations. For each of the combinations of positive selection parameters shown in Table 2, I ran *polyDFE* on uSFS data from 100 bootstrap replicates. When modelling the full uSFS (i.e. with divergence), *polyDFE* identified models containing positive selection consistently for all but one (*p*_*a*_ = 0.0001 and *γ*_*a*_ = 10) of the parameter combinations tested (Table 2). When the DFE was inferred from polymorphism data alone (i.e. without divergence), models containing positive selection were identified less often, particularly when beneficial mutations were rare (*p*_*a*_ = 0.0001; Table 2). Table 2 also shows the proportion of analysis runs for which the gradient of the likelihood exceeded 0.1. The *polyDFE* manual (Tataru & Bataillon, 2019) suggests that gradients >0 indicate that the program has hailed to identify a unique likelihood maximum. When the full uSFS was modelled, the gradient of the likelihood was frequently >0, indicating that the model did not converge on a unique optimum. When modelling the uSFS without divergence, *polyDFE* reported gradients <0.01 for a large proportion of replicate analyses (Table 2).

Figures 3A and 3B show the parameters of positive selection estimated by analysis of uSFS from simulated datasets. I found that when simulated beneficial mutations were mildly advantageous (*γ*_*a*_ = 10) but relatively frequent (*p*_*a*_ = 0.01), both *γ*_*a*_ and *p*_*a*_ were estimated accurately regardless of whether divergence was modelled or not (Figures 3A-B). This finding is consistent with both Schneider et al., (2011) and Tataru et al., (2017). When *p*_*a*_ = 0.01 and *γ*_*a*_ > 10, the analysis of the uSFS with or without divergence yielded very similar parameter estimates, but in both cases, the strength of positive selection seemed to be positively correlated with the estimated *p*_*a*_ (Figure 3). In all cases, when beneficial mutations had *γ*_*a*_ ≥ 50, neither *γ*_*a*_ nor *p*_*a*_ were accurately estimated (Figure 3).

**Figure 3.**
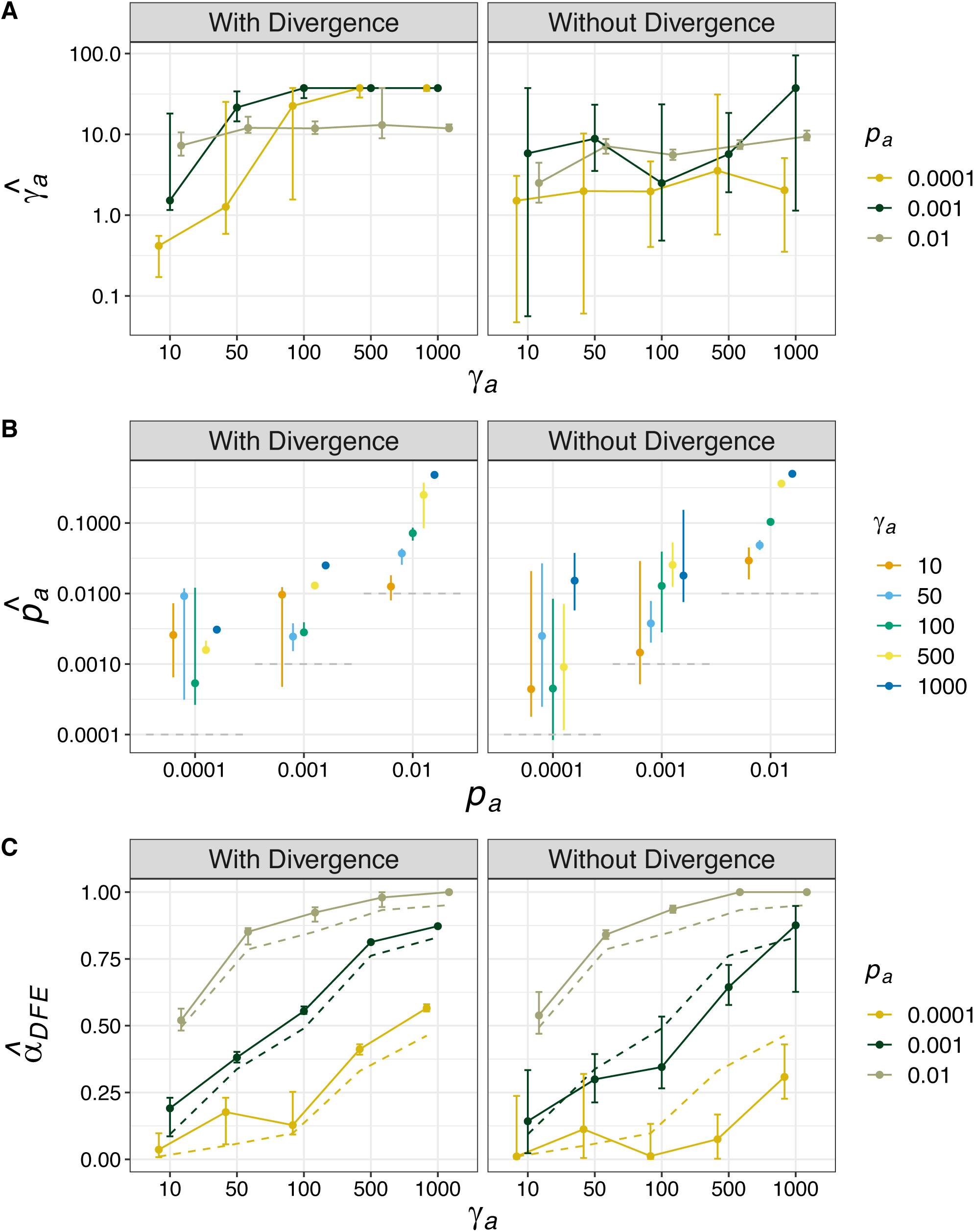
Estimates of the parameters of advantageous mutations and the proportion of adaptive substitutions they imply from simulated datasets. A) *γ*_*a*_ is the inferred selective effect of a new advantageous mutation; B) *p*_*a*_ is the proportion of new mutations that are beneficial, the horizontal dashed grey lines indicate the simulated values in each case; C) *α*_DFE_ is the proportion of adaptive substitutions expected under the inferred DFE, the dashed lines indicate *α*_*Obs*_, the proportion of adaptive substitutions observed in the simulated datasets. Error bars indicate the 95% range of 100 bootstrap replicates.

Tataru et al., (2017) pointed out that, if one had an estimate of the full DFE (i.e. with divergence), the proportion of adaptive substitutions could be obtained by taking the ratio of the fixation probability for a new beneficial mutation over the fixation probability for a random mutation integrating over the full DFE (Equation 10; Tataru et al., 2017). The proportion of adaptive substitutions obtained in this way is denoted *α*_*DFE*_. When modelling the full uSFS, *α*_*DFE*_ was estimated with high accuracy, but with a slight upward bias (Figure 3C). When the DFE was inferred without divergence *α*_*DFE*_ was underestimated when beneficial mutations were strongly selected and rare (Figure 3).

In the presence of infrequent, strongly beneficial mutations the parameters of the DFE for deleterious mutations estimated by *polyDFE* were very accurate (Figure S2). Estimates of the DFE for harmful mutations were less accurate when beneficial mutations occurred with *p*_*a*_ ≥ 0.001 and *γ*_*a*_ ≥ 100. This is presumably because in such cases recurrent selective sweeps eliminate a large amount of neutral diversity and distort the distribution of standing genetic variation at nonsynonymous sites. However, as stated above, the parameter range where the DFE for harmful mutations was poorly estimated in this study may not be biologically relevant.

### Model Identifiability

It is very difficult to tease apart the parameters of positive selection from the uSFS by maximum likelihood. Figure 4 shows the likelihood surface for the three sets of positive selection parameters that satisfy the condition *γ*_*a*_*p*_*a*_ = 0.1. The proportion of adaptive substitutions is largely determined by the product *γ*_*a*_*p*_*a*_ (Kimura & Ohta, 1971) and, as expected, the three parameter combinations shown in Figure 4 all exhibit a similar *α*_*Obs*_ (Figure 1A). However, the extent by which neutral genetic diversity is reduced and the number of segregating advantageous mutations differ substantially across the three parameter combinations (Figure 1). The top row of panels in Figure 4 shows that when modelling the full uSFS, the likelihood surface closely tracks the relation *γ*_*a*_*p*_*a*_ = 0.1. Focussing on the top panel in Figure 4A, the maximum likelihood estimates (MLEs) of the positive selection parameters (the red dot) are far from the true parameter values (indicated by the plus sign), but the MLEs obtained satisfy *γ*_*a*_*p*_*a*_ = 0.1. The ridge in the likelihood surface observed when modelling the full uSFS was described by both Schneider et al., (2011) and Tataru et al., (2017). It comes about because between-species divergence carries information about *α*, and *α* is proportional to *γ*_*a*_*p*_*a*_.

**Figure 4.**
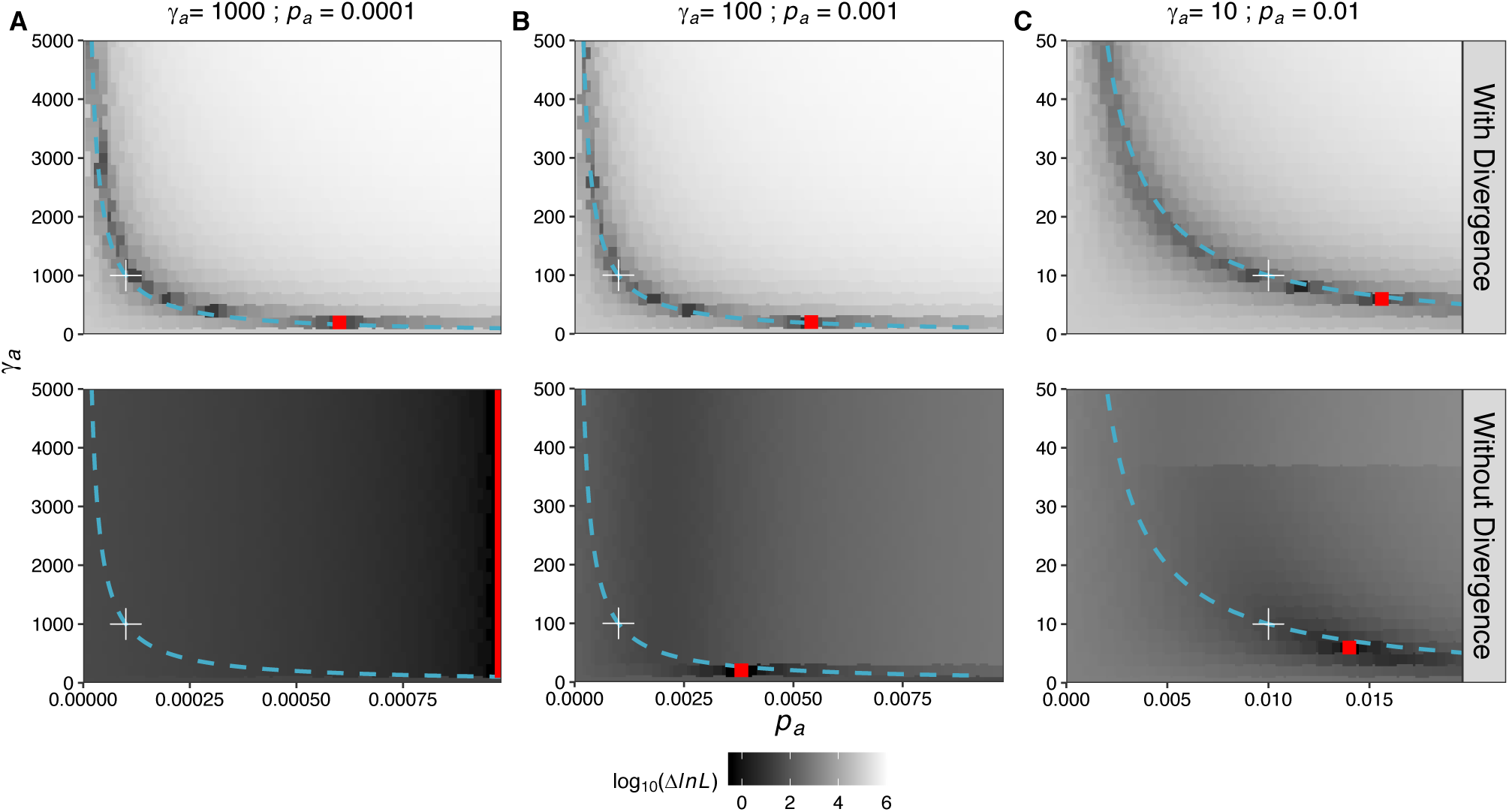
The likelihood surface for the *γ*_*a*_ and *p*_*a*_ parameters for three simulated datasets. Hue indicates differences in log likelihood between a particular parameter combination and the best-fitting model. Best fitting models are indicated by red points and the true parameters are given above the plots and indicated by the white plus signs on the likelihood surface. The relation *γ*_*a*_*p*_*a*_ = 0.1 is shown as a turquoise line and is constant across the three datasets shown.

Inferring the parameters of the DFE from polymorphism alone avoids the assumption of an invariant DFE, but when doing so it may be difficult to distinguish competing models. Indeed, across the three parameter combinations shown, values close to the truth were only obtained from simulated data when *γ*_*a*_ = 10 and *p*_*a*_ = 0.01 (bottom panel Figure 4C). In the case of *γ*_*a*_ = 1000 and *p*_*a*_ = 0.0001, the likelihood surface about the true parameters was very flat (Figure 4A). Increasing the *p*_*a*_ parameter increased likelihood for all strengths of selection, so that the MLEs shown in Figure 4A are simply the values with the highest *p*_*a*_ in the range tested (the vertical red line in Figure 4A). When *γ*_*a*_ = 100 and *p*_*a*_ = 0.001, the likelihood surface about the estimates was steep, but the selection parameters identified by maximum likelihood were incorrect (Figure 4B).

## Discussion

In this study, I analysed simulated datasets modelling a range of positive selection parameter combinations. I found that estimates of positive selection parameters obtained by analysis of the uSFS were only accurate when beneficial mutations had *γ*_*a*_ ≤ 50, under stronger selection the individual parameters of positive selection were not accurately estimated (Figure 3). This is not particularly surprising and is consistent with verbal arguments made in published studies (Booker & Keightley, 2018; Campos et al., 2017). However, it is troubling that when beneficial mutations are strongly selected and rare, the uSFS may often indicate a significant signal of positive selection, but erroneous parameter estimates are obtained. If one were to analyse an empirical dataset and estimate parameters of positive selection of the order *γ*_a_ ∼ 10 and *p*_*a*_ ∼ 0.01, it would be difficult to know whether those were reflective of the true underlying parameters or an artefact of strong selection.

On the basis of this study, it seems that researchers should treat parameters of positive selection obtained by analysis of the uSFS with caution. The expected uSFS for advantageous mutations is very similar when DFE models share the same *p*_*a*_ parameter, and in such cases differing models can only be distinguished by the density of high frequency derived variants (Figure 2). Polarization error when estimating the uSFS can generate an excess in the number of high frequency variants (Keightley & Jackson, 2018), so may generate a spurious signal of strong positive selection. Analysis methods have been proposed which attempt to estimate the rate of polarisation error when modelling the uSFS (Barton & Zeng, 2018; Tataru et al., 2017), but further study is required to determine whether such methods reduce the signal of positive selection in uSFS-based analyses. However, accounting for positive selection when analysing the uSFS yielded robust estimates of the DFE for harmful mutations across the simulated datasets (Figure S2), although I only examined a single DFE for harmful mutations in this study. Tataru et al., (2017) showed that *polyDFE* accurately recovered the parameters of a range of DFE models if positive selection is accounted for.

Estimates of *α* based on analysis of the uSFS may be biased when beneficial mutations are strongly selected and infrequent. Calculating *α* using the rearranged MK-test makes the problematic assumption that the DFE has remained invariant in the time since the focal species began to diverge from the outgroup (Tataru et al., 2017). However, Tataru et al., (2017) pointed out that one can avoid that assumption if *α*_DFE_ is calculated from a DFE estimated without divergence data. In this study, estimates of *α*_*DFE*_ obtained when the full uSFS was analysed were very precise, but with a slight upward bias (Figure 3). When simulated beneficial mutations were strongly selected and rare, the parameters inferred using polymorphism data alone (i.e. without divergence) yielded spurious estimates of *α*_*DFE*_ (Figure 3). When analysing datasets from real populations, *α*_DFE_ may not capture the contribution that strongly beneficial mutations make to molecular evolution. This may make it difficult to contrast *α*_DFE_ between species with large differences in *N*_*e*_, because the number of segregating advantageous mutations and thus ability to accurately estimate selection parameters will depend on the population size.

The nature of the distribution of fitness effects for natural populations is largely unknown. In this study, I analysed the uSFS data under the exact DFE model that had been simulated (i.e. a gamma distribution of deleterious mutational effects plus a discrete class of beneficial effects). However, when analysing empirical data, researchers have to make assumptions about the probability distribution that best describes the DFE of the focal population. A gamma distribution is often assumed for deleterious mutations as it is flexible and is described by only two parameters (Eyre-Walker & Keightley, 2007). However, when analysing real data, one may bias their analyses by strictly adhering to one particular family of probability distributions (Kousathanas & Keightley, 2013). In practice, model averaging provides a way to estimate key features of the DFE while remaining agnostic to the exact shape that the distribution should take (Tataru & Bataillon, 2020). However, if there is bias in the parameter estimates that are obtained across the models that one tests, as is the case for strongly beneficial mutations, a biased average would result.

The simulations I performed in this study generated the ideal dataset for estimating parameters of selection from the uSFS. I simulated 21Mbp of coding sites in which genotypes and whether sites were selected or not was unambiguously known. When analysing real data this is not the case and researchers often have to filter a large proportion of sites out of their analyses or choose to analyse a subset of genes that have orthology with outgroups or other biological properties of interest. Even with perfect knowledge, strongly beneficial mutations only represented a small proportion of the standing genetic variation at nonsynonymous sites (Figure 1, S1). In addition, the populations I simulated were randomly mating and had constant sizes over time. The results I present in this study suggest that even with perfect knowledge of a population that adheres to the assumptions of a Wright-Fisher model, it is inherently difficult to infer the parameters of strongly beneficial mutations from the uSFS, particularly so when beneficial mutations occur infrequently.

### Estimating parameters of positive selection from the uSFS versus estimates from patterns of diversity

As discussed above, studies based on analysis of the uSFS and those based on selective sweep models have yielded vastly different estimates of the parameters of positive selection. Patterns of neutral genetic diversity in both humans and wild mice cannot be explained by the effects of background selection alone, and in both species it has been suggested that strongly beneficial mutations are required to explain the observed patterns (Booker & Keightley, 2018; Nam et al., 2017). In the case of wild house mice, positive selection parameters obtained by analysis of the uSFS do not explain dips in nucleotide diversity around functional elements (Booker & Keightley, 2018). Recently, Castellano et al. (2019) analysed the uSFS for nonsynonymous sites in great ape species but did not find significant evidence for positive selection. In their dataset, Castellano et al. (2019) had at least 8 haploid genome sequences for each of great ape species they analysed, and they argued that they were underpowered to detect positive selection on the basis of the uSFS. In this study, I analysed datasets of 20 diploid individuals and found that it was very difficult to accurately capture positive selection parameters. Increasing the number of sampled individuals even further may increase the power to estimate the strength of positive selection, but this study suggests that the increase in power will depend on the underlying DFE. When *p*_*a*_ is small, the expected number of advantageous mutations present in the uSFS for 200 diploids is less than 10 for most frequency classes when14Mbp of nonsynonymous sites have been used to construct the uSFS (Figure S3). Indeed, Figure S3 shows that even with very large sample sizes, the expected uSFS for beneficial mutations are very similar and may only be distinguished on the basis of a small number of high frequency derived alleles. Thus, it may be that the uSFS is inherently limited in the information it carries on the DFE for beneficial mutations so other sources of information may have to be used to accurately recover parameters.

In this study, I modelled beneficial mutations using a discrete class of selection coefficients when, in reality, there is likely a continuous distribution of fitness effects. Indeed, studies in both humans and *D. melanogaster* have found evidence for a bimodal distribution containing both strongly and weakly beneficial mutations contributing to adaptive evolution using methods which incorporate linkage information but do not explicitly estimate selection parameters (Elyashiv et al., 2016; Uricchio et al., 2019). There are currently no methods that estimate the DFE using an analytical expression for the uSFS expected under the combined effects of BGS and sweeps. Rather, nuisance parameters or demographic models are used to correct for the contribution that selection at linked sites may make to the shape of the SFS (Eyre-Walker, Woolfit, & Phelps, 2006; Galtier, 2016; Tataru et al., 2017). However, as this study shows, the parameters of positive selection are not reliably estimated when analysing the uSFS alone. A way forward may be in using computational approaches to make use of all of the available data, while not necessitating an expression for the uSFS expected under the combined effects of BGS, sweeps, population size change and direct selection. An advance in this direction has recently been made by Uricchio et al., (2019) who developed an ABC method for estimating *α* which makes use of the distortions to the uSFS generated by BGS and sweeps. By applying their method to data from humans, Uricchio et al., (2019) found that *α* = 0.13 for nonsynonymous sites, 72% of which was generated by mildly beneficial mutations and 28% by strongly beneficial mutations. However, the computational approach developed by Uricchio et al., (2019) could readily be extended to model an arbitrarily complex DFE for beneficial mutations. Their methods could be implemented in a machine-learning context, with training data generated by forward-simulations that capture confounding factors such as population structure and population size change as well as the effects of selection at linked sites.

## Acknowledgements

I wish to extend gratitude to Peter Keightley, Michael Whitlock and Sam Yeaman for valuable advice and mentorship. Thanks to Thomas Bataillon, Sam Yeaman and Peter Keightley for comments on previous versions of the manuscript. Thanks to Paula Tataru and Thomas Bataillon for help with *polyDFE*. Thanks to two anonymous reviewers for constructive feedback.

## Supplementary Material

**Figure S1.**
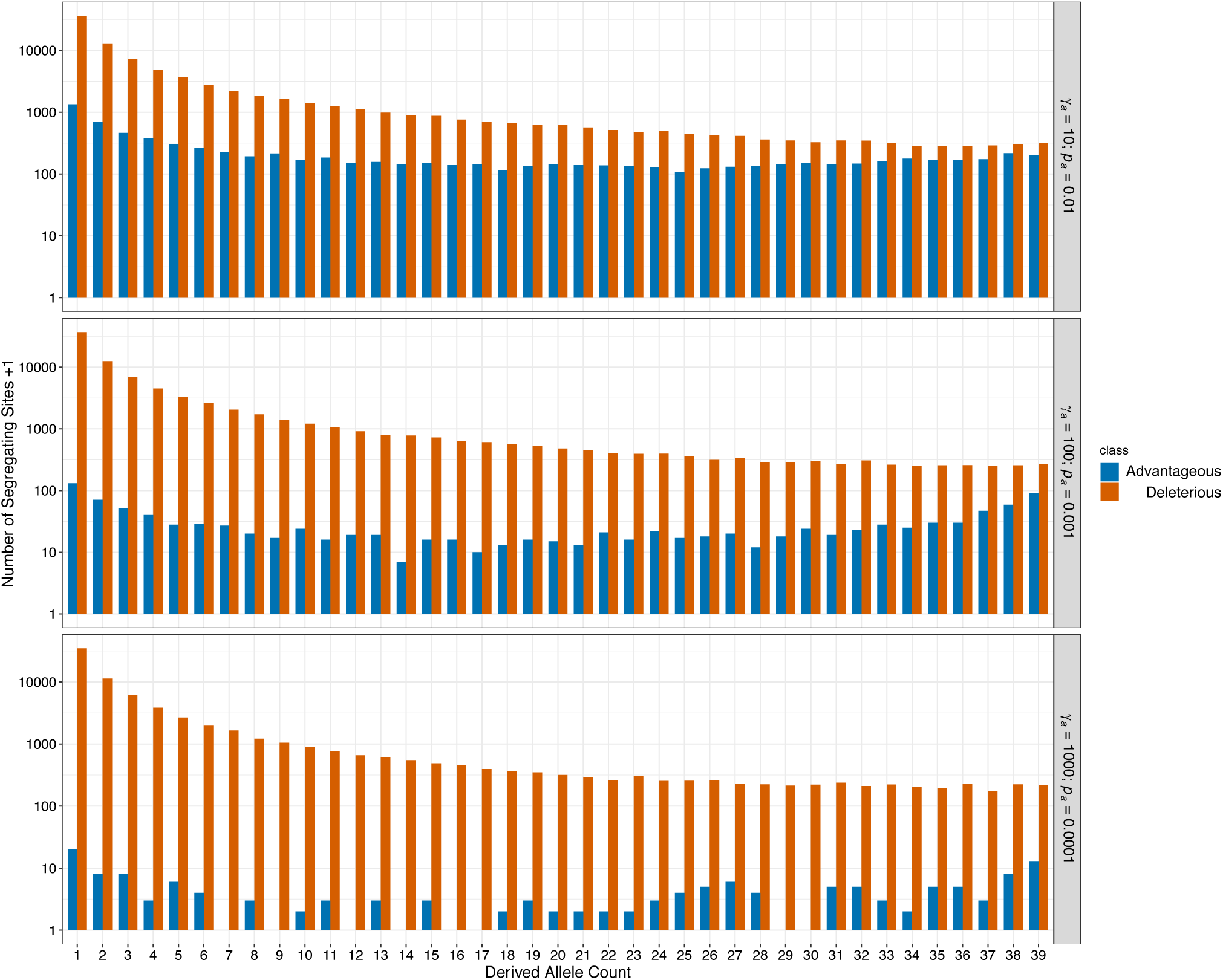
The observed uSFS for nonsynonymous sites for three sets of positive selection parameters. The distribution of deleterious mutations is shown in orange and the distribution of advantageous mutations is shown in blue. For the purposes of visualising the data on a log scale, the number of segregating sites is shown +1.

**Figure S2.**
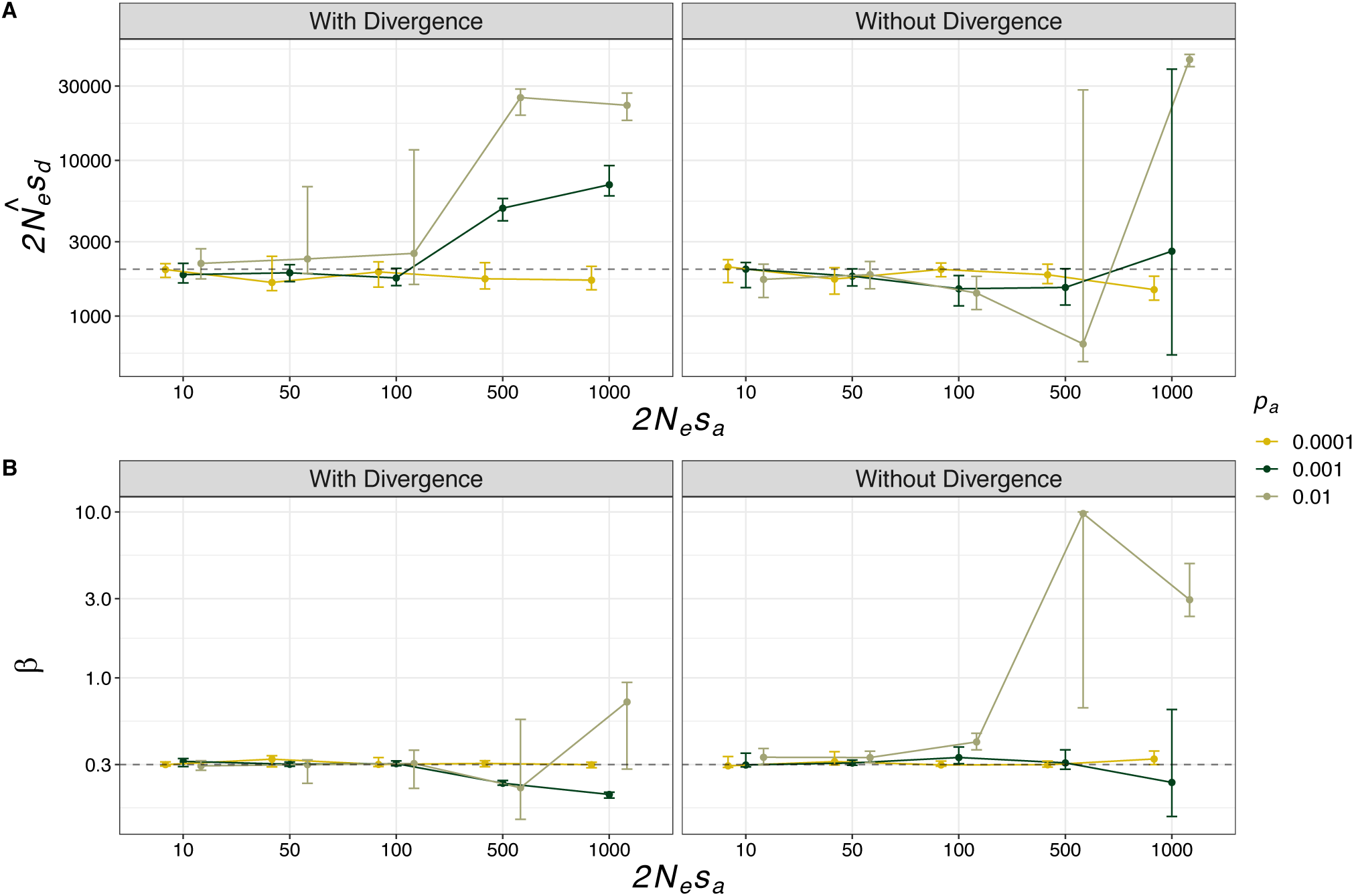
Parameter estimates for the DFE for deleterious mutations obtained from simulated datasets. A) the mean effect of a deleterious mutation and b) the shape parameter of the gamma distribution. Error bars indicate the 95% range of 100 bootstrap replicates.

**Figure S3.**
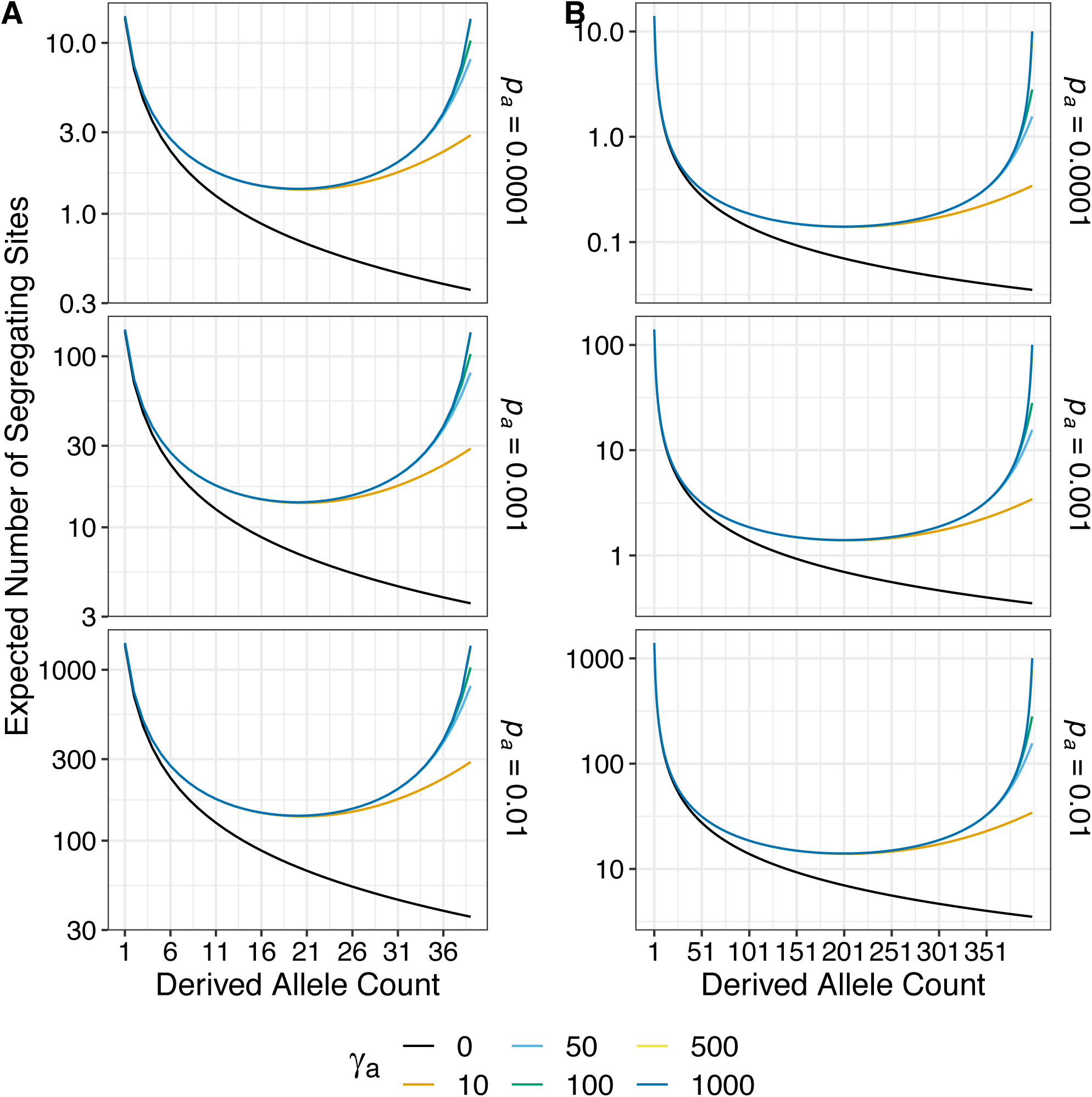
The expected uSFS for beneficial alleles. Panel A shows the expected uSFS for a sample of 20 diploid individuals, and panel B shows the uSFS for 200 diploid individuals.

## Notes

#### Summary of Updates

This version of the manuscript has been revised in response to a round of peer-review.

